# Chromosome-scale genome assembly of the North American invasive Asteraceae, *Tanacetum vulgare* (common tansy)

**DOI:** 10.64898/2026.09.30.755630

**Authors:** Dimiru Tadesse, Adrian E. Platts, Lori Croghan, Alan G. Smith, Neil O. Anderson, Roger L. Becker, Patrick P. Edger, Matthew A. Tancos

## Abstract

Common tansy (*Tanacetum vulgare*), a significant agricultural invasive species, threatens forage productivity, livestock carrying capacity, and plant biodiversity across North American agroecosystems. Despite being a close relative to the ornamental Asteraceae *Chrysanthemum, T. vulgare* genomic studies are severely lacking, but remain critical to the development of novel weed management tools and the investigation of plant invasion evolution. In this study, we performed the assembly and annotation of the diploid genome (2*n*=2*x*=18) of an invasive North American *T. vulgare* genotype using long-read PacBio HiFi and Iso-Seq sequencing technology. The assembled haplotype genome comprised 9 pseudomolecules with a total size of ∼4.8 Gb and a scaffold N50 of 546 Mb. In addition, both the plastid and mitochondrial genome sequences were assembled. BUSCO analysis revealed a high overall genome completeness with 69,343 predicted protein-coding genes (AED<0.6: 98%). A high-quality North American invasive reference *T. vulgare* genome, complementing a recent one assembled for a native European accession, is critical for the investigation of plant invasion biology, genome evolution, and the future development of novel genetic biocontrol weed management technologies.

## Background & summary

Invasive alien species are a US$1.288 trillion financial strain to the global economy, negatively impacting agriculture, infrastructure, and public health ^1^. In the United States invasive plants are estimated to cause US$35 billion in agriculture-related losses annually, representing significant costs to growers, ranchers, foresters, and land managers ^2^. Weed management strategies outside of highly mechanized cropping systems are difficult due to the complex, heterogenous terrain of rangelands, forests, and natural systems, spanning millions of acres and supporting diverse beneficial and native plant species ^3^. Current chemical, mechanical, and cultural management practices often are not cost-effective or practical for weed species infesting vast, complex, natural landscapes. In contrast, biological control is a management strategy well-suited for these complex ecosystems, but many invasive target species currently lack suitable, federally approved agents ^4^. To help address these challenges, next generation weed management strategies need to leverage genomics and gene-editing applications to develop novel genetic biocontrol tools ^4^. Development of high-quality genomic resources for high-impact invasive species is a critical step to advance species-specific genetic biocontrol tools, such as a weed-suppressive gene drive^5^.

Common tansy (*Tanacetum vulgare* L, Asteraceae) is an invasive perennial herb native to temperate regions of Eurasia and was introduced into North America during the 17^th^ century by European settlers as a funerary, medicinal, and ornamental herb ^6,7^. Earliest records document its arrival at the New England Plymouth colony by John Winthrop Jr. in 1631 ^6^. Today, *T. vulgare* is naturalized across the US and Canada with dense populations primarily infesting ruderal and natural environments of the northern US and southern Canada ^7,8^. *Tanacetum vulgare* primarily spreads sexually (localized spread is enhanced by clonal propagation via rhizomes) as it produces abundant outcrossing flowerheads (>100/plant) and up to 50,000 seeds/plant, outcompeting beneficial and native forage plants, impeding timberland regeneration, and threatening livestock health and milk production due to the production of toxic secondary metabolites ^5,7,9-11^. While the production of diverse terpenes, flavonoids, and essential oils negatively impact livestock and herbivorous insects, these chemistries also make *T. vulgare* an attractive target for pharmacological and insecticidal investigations ^12,13^. The successful invasion of North America, deterrence of endemic predators, adaptability to diverse environments, high fecundity, continued use as an ornamental (sold on the cut flower market), and threat to US agriculture made *T. vulgare* an ideal invasive species to investigate plant invasion genomics and novel genetic biocontrol tools.

The purpose of this study was to generate a high-quality annotated genome assembly for invasive *T. vulgare*. Here, we present a chromosome-level genome assembly and annotation of a North American invasive *T. vulgare* genotype, produced using PacBio HiFi long-read sequencing and Iso-Seq transcript data. The assembly was scaffolded with a European *T. vulgare* accession collected in the United Kingdom (GCA_964264315.1) using RagTag ^14,15^, resulting in nine pseudochromosomes and an estimated haploid genome size of approximately 4.8 Gb. The availability of genomic resources from both a European genotype representing the species’ native range and a North American genotype from its invaded range provides an important foundation for comparative genomic analyses. Together, these assemblies will enable investigation of genetic variation, genome structure, and evolutionary processes associated with invasion success. More broadly, this resource will support studies of plant invasion biology, adaptation, and genome evolution, while also facilitating the development and assessment of genetic biocontrol approaches, such as a weed-suppressive gene drive for managing invasive *T. vulgare* populations^5^.

## Materials and methods

### Plant material and extractions

*Tanacetum vulgare* seed was field-collected and pooled from multiple Minnesota, US, infestations and germinated under ambient greenhouse conditions (∼20-25°C) with natural light in a commercial soilless mix (Sunshine Mix #1, Sun Gro, Agawam, MA, USA). A single emergent seedling was selected and transferred into a 4-inch pot (T.O. Plastics, Inc., Clearwater, MN, USA) and fertilized using a slow-release fertilizer per manufacturer instructions (Osmocote 14-14-14, The Scotts Company, Marysville, OH, USA). All plant material used for long-read sequencing was collected from this individual *T. vulgare* genotype (Figure 1). Fresh, emergent leaf tissue was collected and flash-frozen in liquid nitrogen, briefly macerated in a TissueLyser II (Qiagen, Germantown, MD, USA), and processed using the Promega Wizard HMW extraction kit (Promega Corporation, Madison, WI, USA). Total RNA was isolated from flash-frozen tissue collected from the root, stem, leaves, and flowers, and extracted using the Qiagen RNeasy Kit (Qiagen, Germantown, MD, USA). Nucleic acid was shipped overnight on ice (DNA) or dry ice (RNA) to the Delaware Biotechnology Institute at the University of Delaware DNA Sequencing & Genotyping Center for PacBio HiFi and IsoSeq workflows (Pacific Biosciences, CA, USA).

**Figure 1.).**
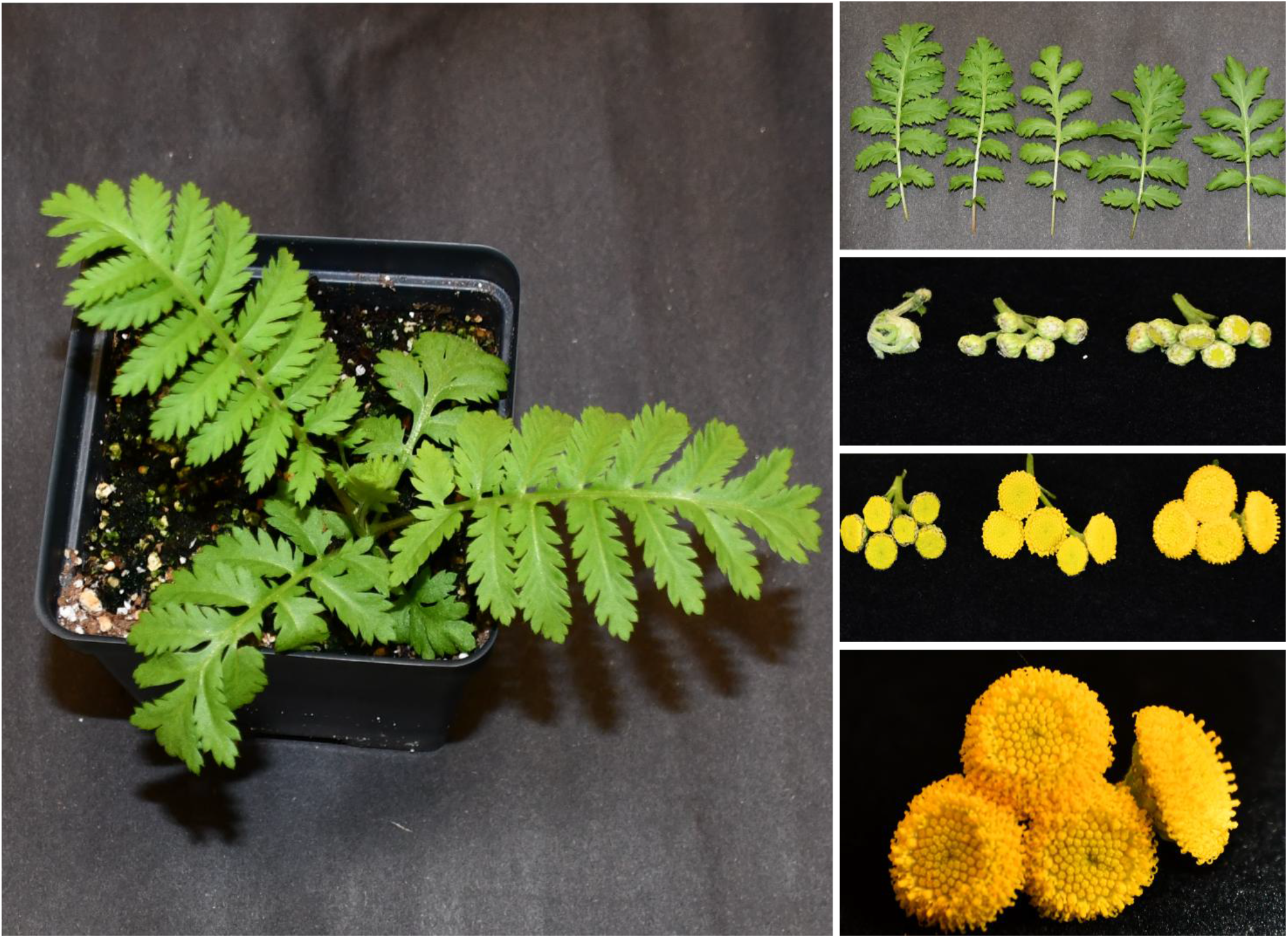
Morphological characteristics of North American *Tanacetum vulgare*. Tansy plant grown under ambient greenhouse conditions (approximately 20–25°C) with natural light in a commercial soilless mix at approximately 3-weeks-old. Images illustrate leaf morphology and floral developmental stages.

### DNA/RNA sample integrity analyses

DNA quality was assessed using Femto Pulse (Agilent, CA, USA), Nanodrop and Qubit fluorometric quantification to assess DNA quality and the presence of long, intact fragments suitable for HiFi library preparation. RNA integrity from root, stem, leaf, and flower tissue samples were evaluated using a capillary electrophoresis Fragment Analyzer (Agilent, CA, USA), ensuring high-quality RNA samples with high >7 RIN values ^16^.

### Library preparation and sequencing

PacBio HiFi libraries were prepared following manufacturer protocols for high-fidelity genome sequencing using SMRTbell library kit 3.0. The library was enriched with fragments greater than 8kb by size selection using Blue Pippin (Sage Science, MA, USA). For transcriptome sequencing, IsoSeq libraries were generated independently for each tissue, employing full-length cDNA synthesis and SMRTbell library construction optimized for long-read RNA sequencing.

The genomic DNA library was sequenced on the Sequel IIe instrument using six SMRT cells with 30 hours of movie time. IsoSeq libraries for each tissue type were sequenced individually on Sequel IIe instrument with 24 hours movie sequencing. IsoSeq data were analyzed using PacBio Iso-seq 3 flow producing full-length non-concatemer reads with polyA tail. A total of ∼338Gb of data was generated equating to 70.6x coverage (Table 1).

**Table 1).** Sequencing data statistics for *Tanacetum vulgare*.

| Data source | Read count | Total data (Gb) | Genome coverage (X) |
| --- | --- | --- | --- |
| PacBio HiFi | 13015088 | 338.887 | 70.6 |
| Iso-Seq (root) | 233549 | 6.273 | — |
| Iso-Seq (stem) | 110577 | 2.931 | — |
| Iso-Seq (leaf) | 188668 | 5.071 | — |
| Iso-Seq (flower) | 128163 | 3.453 | — |

### Genome survey and heterozygosity

The K-mer frequencies were quantified from the aggregated PacBio HiFi FASTQ datasets using Meryl v1.4.1 ^17^. K-mers of length 21 were counted using the command: *meryl threads=32 k=21 count All*.*gz output tansy*.*meryl*. The resulting k-mer database was subsequently transformed into a full k-mer frequency histogram without an upper limit using: *meryl histogram tansy*.*meryl > tansy*.*hist*. GenomeScope2 was then used to model genome size from the observed k-mer distribution ^18^. The histogram generated by Meryl served as input to GenomeScope2, using the command: *genomescope2 -i tansy*.*hist -o output_directory -k 21 -p 2*. The model estimated a haploid genome size of approximately 4.5 Gb, which was ∼7% smaller than the final assembled genome size. Upon inspection of the model fit (Figure 2), we noted that GenomeScope2 did not fully accommodate the extended high-frequency tail of the k-mer distribution. This discrepancy likely contributed to the slight underestimation of genome size. Based on the modeled k-mer profile, the heterozygosity rate of the genome was estimated at 1.56%.

**Figure 2).**
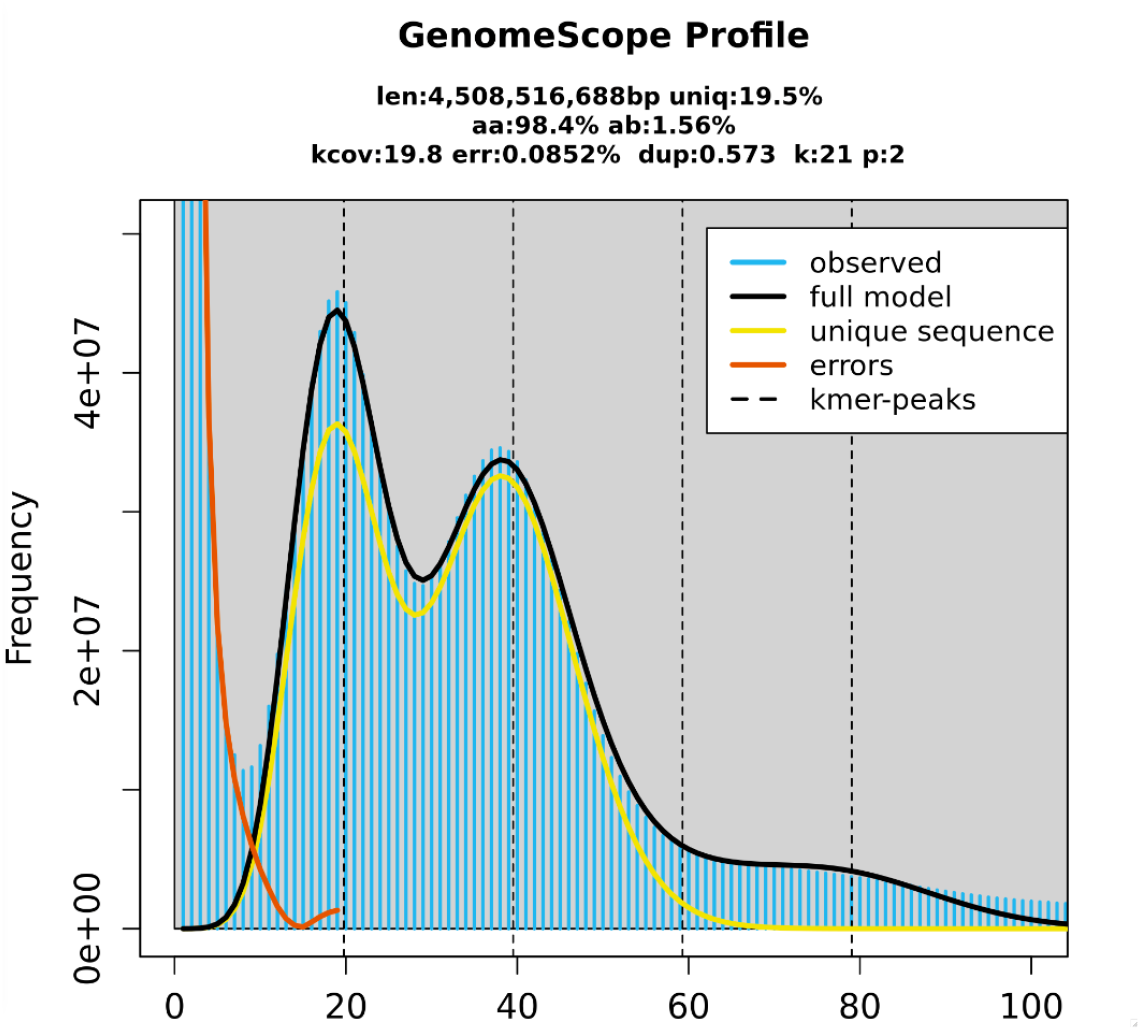
K-mer based genome size and heterozygosity estimation. The GenomeScope profile illustrates the k-mer coverage distribution used to model genome characteristics of *Tanacetum vulgare*. Blue bars represent the observed k-mer frequencies, and the black line shows the fitted GenomeScope model. The yellow curve indicates the predicted unique sequence component, and the red curve highlights sequencing error k-mers. Vertical dashed lines mark the major k-mer peaks associated with distinct coverage modes.

### Genome assembly and scaffolding

The North American *T. vulgare* genome was assembled from six SMRT cells of Pacific Biosciences HiFi data using HiFiAsm ^19^ with default parameters. The initial assembly comprised 5.14 Gb in approximately 4,000 contigs. Nanopore ultra-long sequencing failed to yield usable reads; scaffolding was therefore performed using RagTag ^14^ with a European *T. vulgare* accession collected in the United Kingdom (GCA_964264315.1) as a reference ^15^. RagTag was run sequentially in correct mode (correct -f 10000 --remove-small) followed by scaffold mode, removing contigs shorter than 10 kb. The assembly produced an overall genome size of approximately 4.8 Gb and achieved a scaffold N50 of 546 Mb (Table 2).

**Table 2).** Assembly metrics for the primary and scaffolded *Tanacetum vulgare* genome.

| Item | Scaffolded assembly |  | Primary assembly |  |
| --- | --- | --- | --- | --- |
|  | N | % | N | % |
| Number of scaffolds | 11 |  | 4,308 |  |
| Total size of scaffolds | 4,846,764,676 |  | 5,142,052,626 |  |
| Longest scaffold | 647,928,270 |  | 17,738,146 |  |
| Shortest scaffold | 272,090 |  | 10,574 |  |
| Number of scaffolds > 500 nt | 11 | 100% | 4,308 | 100% |
| Number of scaffolds > 1K nt | 11 | 100% | 4,308 | 100% |
| Number of scaffolds > 10K nt | 11 | 100% | 4,308 | 100% |
| Number of scaffolds > 100K nt | 11 | 100% | 2,819 | 65% |
| Number of scaffolds > 1M nt | 9 | 82% | 1,530 | 36% |
| Mean scaffold size | 440,614,971 |  | 1,193,606 |  |
| Median scaffold size | 538,527,034 |  | 425,518 |  |
| N50 scaffold length | 546,907,052 |  | 3,088,220 |  |
| L50 scaffold count | 5 |  | 491 |  |
| N50 scaffold - NG50 scaffold length difference | 546,907,052 |  | 3,088,220 |  |
| scaffold %A | 32 |  | 32 |  |
| scaffold %C | 18 |  | 18 |  |
| scaffold %G | 18 |  | 18 |  |
| scaffold %T | 32 |  | 32 |  |
| scaffold %N | 0 |  | 0 |  |
| scaffold N nt | 287,200 |  | 0 |  |
| scaffold %non-ACGTN | 0 |  | 0 |  |
| Number of scaffold non-ACGTN nt | 0 |  | 0 |  |
| Percentage of assembly in scaffolded contigs | 100% |  | 0% |  |
| Percentage of assembly in unscaffolded contigs | 0% |  | 100% |  |
| Average number of contigs per scaffold | 262 |  | 1 |  |
| Average length of breaks (20 or more Ns) between contigs | 100 |  | 0 |  |

### Genome annotation

Gene annotation was performed using MAKER ^20^, integrating EST evidence from StringTie ^21^ in long-read mode based on IsoSeq data from leaf, root, stem and flower tissues, protein homology data from *Chrysanthemum indicum* ^22,23^, and Augustus gene models. Additional downstream filtering involved removal of gene models with transposon domains as reported by InterProScan ^24^, merger of gene models showing complementary alignment to the same *C. indicum* protein (DIAMOND ^25^), and rejection of weak maker models with an AED score more than 0.6. The nine pseudomolecules range from 365 to 648 Mb in length and together form a 4.8 Gb genome. They contain a total of 69,343 predicted genes, with individual chromosomes carrying between about 5,113 and 9,684 genes (Table 3).

**Table 3).** Nine pseudochromosomes, length and total number of genes in each chromosome of *Tanacetum vulgare*.

| Pseudochromosome | GCA_964264315.1 Names | Length (Mb) | Genes |
| --- | --- | --- | --- |
| TvChr01 | OZ183338 | 656.02 | 9684 |
| TvChr02 | OZ183339 | 620.00 | 8162 |
| TvChr03 | OZ183340 | 579.29 | 7430 |
| TvChr04 | OZ183341 | 507.52 | 6962 |
| TvChr05 | OZ183342 | 553.74 | 7101 |
| TvChr06 | OZ183343 | 567.53 | 8846 |
| TvChr07 | OZ183344 | 545.25 | 8571 |
| TvChr08 | OZ183345 | 370.28 | 5113 |
| TvChr09 | OZ183346 | 506.74 | 7372 |
| TvChr10 (mitochondria) | OZ183347 | 27.54 | 22 |
| TvChr11 (plastid) | OZ183348 | 64.72 | 80 |
| Total |  | 4.8 Gb | 69,343 |

### Technical validation of the genome assembly

Long-read HiFi data quality was assessed using NanoPlot, confirming high read accuracy and appropriate read-length distributions (median = 13,332bp). Genome assembly completeness was evaluated using BUSCO (Benchmarking Universal Single-Copy Orthologs) with the eudicotyledons_odb12 lineage dataset and Busco version busco6/6.0.0 ^26^, using the command: *busco -i ragtag*.*scaffold*.*pseudo*.*folded*.*H1*.*fasta -l eudicotyledons_odb12 -m genome -o final_for_paper -c 32 -f*. The resulting assembly showed high completeness, consisting of 2776 (99%) complete BUSCOs, including 2344 (83.6%) complete single-copy and 432 (15.4%) complete duplicated BUSCOs, with 6 (0.2%) fragmented and 23 (0.8%) missing BUSCOs (Table 4). Transcriptome completeness was similarly assessed using BUSCO with the eudicotyledons_odb12 database, run using the command: *busco -i final*.*pep*.*fa -l eudicotyledons_odb12 -m protein -o tx_for_paper -c 32 -f*. The resulting transcriptome consisted of 2255 (80.4%) complete BUSCOs, including 1933 (68.9%) complete single-copy and 322 (11.5%) complete duplicated BUSCOs, with 207 (7.4%) fragmented and 343 (12.2%) missing BUSCOs (Table 5). This analysis provided an independent validation of transcriptome quality and supported the robustness of the assembled gene space. Additional transcriptome work will continue as further resources become available.

**Table 4).** BUSCO analysis of *Tanacetum vulgare* genome assembly.

| Category | Code | Count | Percentage |
| --- | --- | --- | --- |
| Complete BUSCOs | C | 2776 | 99.0% |
| Complete and single-copy BUSCOs | S | 2344 | 83.6% |
| Complete and duplicated BUSCOs | D | 432 | 15.4% |
| Fragmented BUSCOs | F | 6 | 0.2% |
| Missing BUSCOs | M | 23 | 0.8% |
| Total BUSCOs |  | 2805 | - |

**Table 5).** BUSCO analysis of *Tanacetum vulgare* transcriptome.

| Category | Code | Count | Percentage |
| --- | --- | --- | --- |
| Complete BUSCOs | C | 2255 | 80.4% |
| Complete and single-copy BUSCOs | S | 1933 | 68.9% |
| Complete and duplicated BUSCOs | D | 322 | 11.5% |
| Fragmented BUSCOs | F | 207 | 7.4% |
| Missing BUSCOs | M | 343 | 12.2% |
| Total BUSCOs |  | 2805 | - |

### Functional annotation of invasive North American *T. vulgare* proteome

Functional annotation was performed using a non-redundant protein dataset. Only amino acid sequences with valid open reading frames (ORFs) were retained for similarity searches. Reference protein datasets SwissProt and its supplement TrEMBL ^27^ were used for homology-based functional annotation. Amino acid FASTA files for each reference dataset were downloaded from their respective public repositories and converted into diamond formatted databases with the makedb utility using diamond v2.1.24 ^28^:*diamond makedb --in uniprot_sprot*.*fasta -d swissprot*.*dmnd* and for TrEMBL, *diamond makedb --in uniprot_trembl*.*fasta -d trembl*.*dmnd*. Sequence similarity searches were then performed using diamond v2.1.24 in blastp mode, chosen for its high speed and sensitivity suitable for large scale genome and transcriptome annotation ^28^. Each protein sequence was queried independently against the SwissProt, and TrEMBL databases using the following parameters: E-value threshold (1e−5), maximum target hits (25), substitution matrix (BLOSUM62), gap penalties (default ,11/1), to get a tabular formatted file containing the best scoring alignments. The diamond output files (swissprot.out, trembl.out) contained alignment scores, percentage identity, alignment length, query coverage, and database identifiers. Unique proteins with only one stop codon were screened and *Tansy_proteins*.*noStop*.*fasta* was generated. These results were retained for integration with domain based functional annotation (InterProScan), and motif identification using Pfam (protein family database) and hmmscan (Profile Hidden Markov Model-based scan). To identify protein domains and functional motifs, protein structural and functional features were annotated using InterProScan (v5.x) ^24^, which integrates multiple domain and motif databases including Pfam, SMART (Simple Modular Architecture Research Tool), CDD (conserved domain database), PROSITE (protein site), Panther (protein analysis through evolutionary relationships), TIGRFAMs (The Institute for Genomic Research Protein Families), and Gene3D. All predicted protein sequences were analyzed in standalone mode with the following command: *interproscan*.*sh -i Tansy_proteins*.*noStop*.*fasta --formats TSV,XML,GFF3 --goterms -- pathways --iprlookup --cpu 32 --output-dir iprscan_out*. Domain hits with full length coverage across Pfam and SMART families were used to refine gene function predictions and support downstream classification and visualization. Protein domain annotation was performed using HMMER v3.4 ^29^ against the Pfam-A database. Domain detection was performed using HMMER (hmmscan), and detailed per domain hit information was captured using the --domtblout output format:

*hmmscan --domtblout pfam*.*domtblout Pfam-A*.*hmm Tansy_proteins*.*noStop*.*fasta*. The resulting pfam.domtblout file contains domain level information for each protein, including the Pfam identifier, domain coordinates on the query sequence, and scoring statistics. To extract domain architectures, the domtblout file was parsed to retain only protein identifiers, Pfam accession names, and domain start/end coordinates. All domain hits were sorted by their start positions and concatenated to produce ordered domain architecture strings for each protein. Unique domain architectures were then quantified across the proteome. The frequency of each Pfam domain across the entire annotated protein set was quantified by counting domain occurrences per protein. Domains were ranked by total abundance, and the 20 most frequently represented Pfams were selected for visualization (Figure 3a). For each protein, Pfam domains were grouped to define its domain architecture. Domain combinations were tabulated genome-wide to identify recurring pairings and multidomain patterns (Figure 3b).

**Figure 3).**
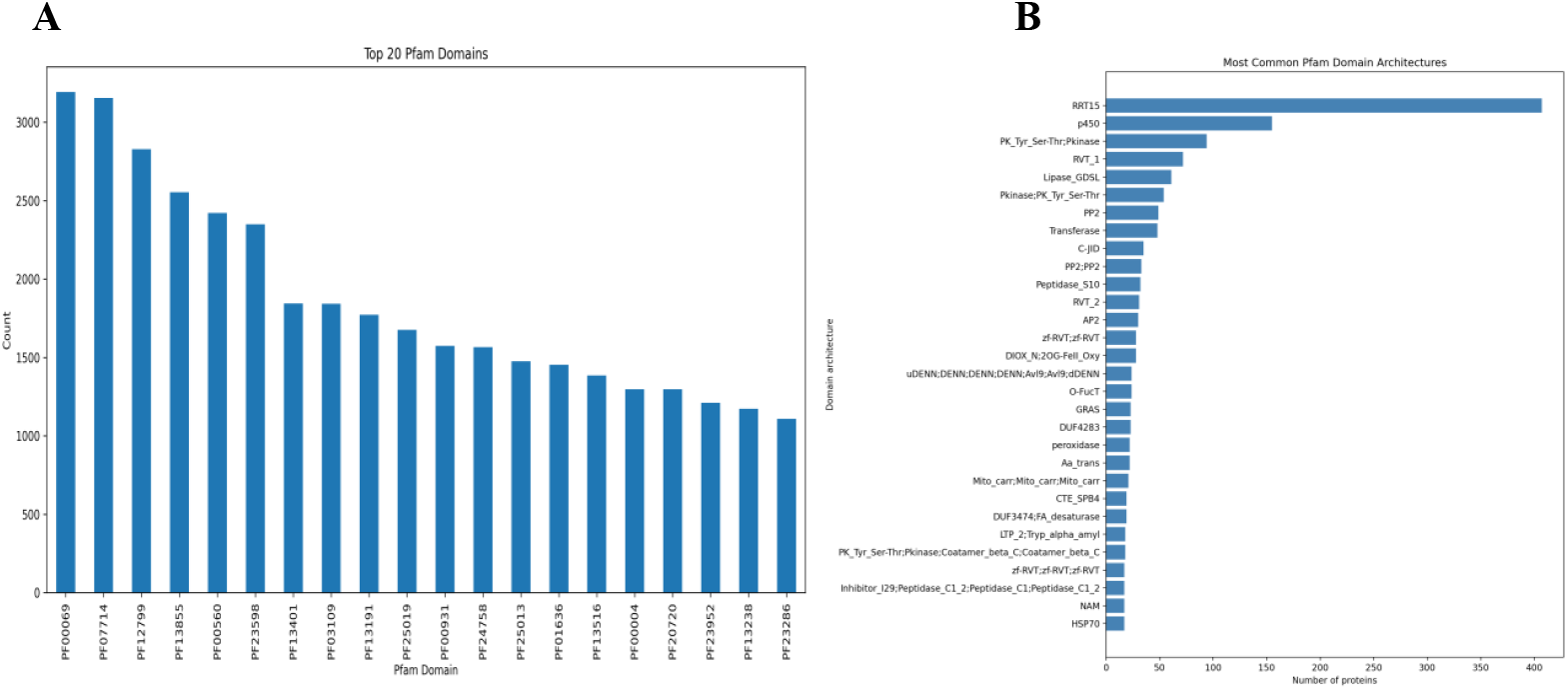
Functional annotation of invasive North American *Tanacetum vulgare* proteome. (A) shows twenty most abundant Pfam protein domains detected across the annotated genome. Domain counts reflect the number of predicted proteins containing each Pfam entry, highlighting the most frequently occurring functional domains within the *T. vulgare* proteome. (B) shows the most frequently occurring Pfam domain architectures identified in the genome annotation. Each architecture represents a unique combination of Pfam domains found within individual predicted proteins, highlighting how domains are organized and reused across the proteome. Higher counts indicate domain arrangements that are widely conserved or expanded in the genome, often corresponding to essential biological processes such as transcriptional regulation, enzymatic metabolism, signaling, and structural maintenance.

### Genome-wide analysis of secondary metabolite biosynthetic gene clusters in *T. vulgare*

Secondary metabolite biosynthetic analyses were performed with the plantiSMASH v2.0.4 standalone program ^30^ using USDA’s SCINet HPC platform following the documented command-line workflow for FASTA + GFF3 inputs, which requires matching sequence identifiers between the FASTA headers and the GFF3 seqid column. FASTA–GFF3 name alignment was verified and used the plantiSMASH plant taxon preset with CDS phase awareness (--use_phase) and no contig length limit (--limit -1). The GFF3 annotation was standardized prior to analysis to conform to the GFF3 specification. Formatting and structural issues were corrected via Genome Tools (gt gff3validator for strict validation; gt gff3 -tidy -sort -checkids -retain ids for normalization), followed by removal of blank lines to satisfy strict parsers. PlantiSMASH was run using the command: *plantismash --taxon plants --use_phase --limit -1 --cpus 32 -- disable_subgroup --disable_treesvg --gff3 tansy*.*clean9*.*gff3 tansy_corrected*.*fasta --outputfolder result_tansy_no_subgroup --verbose*, disabling the subgroup finder and tree rendering to focus on core cluster detection. The completed run produced an interactive HTML report with per-cluster SVG schematics (svg/geneclusters) and excel summaries. Across contigs OZ183338 - OZ183346, there are 31 predicted specialized secondary biosynthesis clusters each containing 4 - 54 genes. While majority of the clusters are dominated by cyclopeptide and alkaloids, recurring core domains such as UGTs/transferases, 2OG-Fe(II) oxygenases, P450s, and Bet_v_1/PR-10 suggests glycosylation, oxidation, and cyclization tailoring steps (Table 6).

**Table 6).**
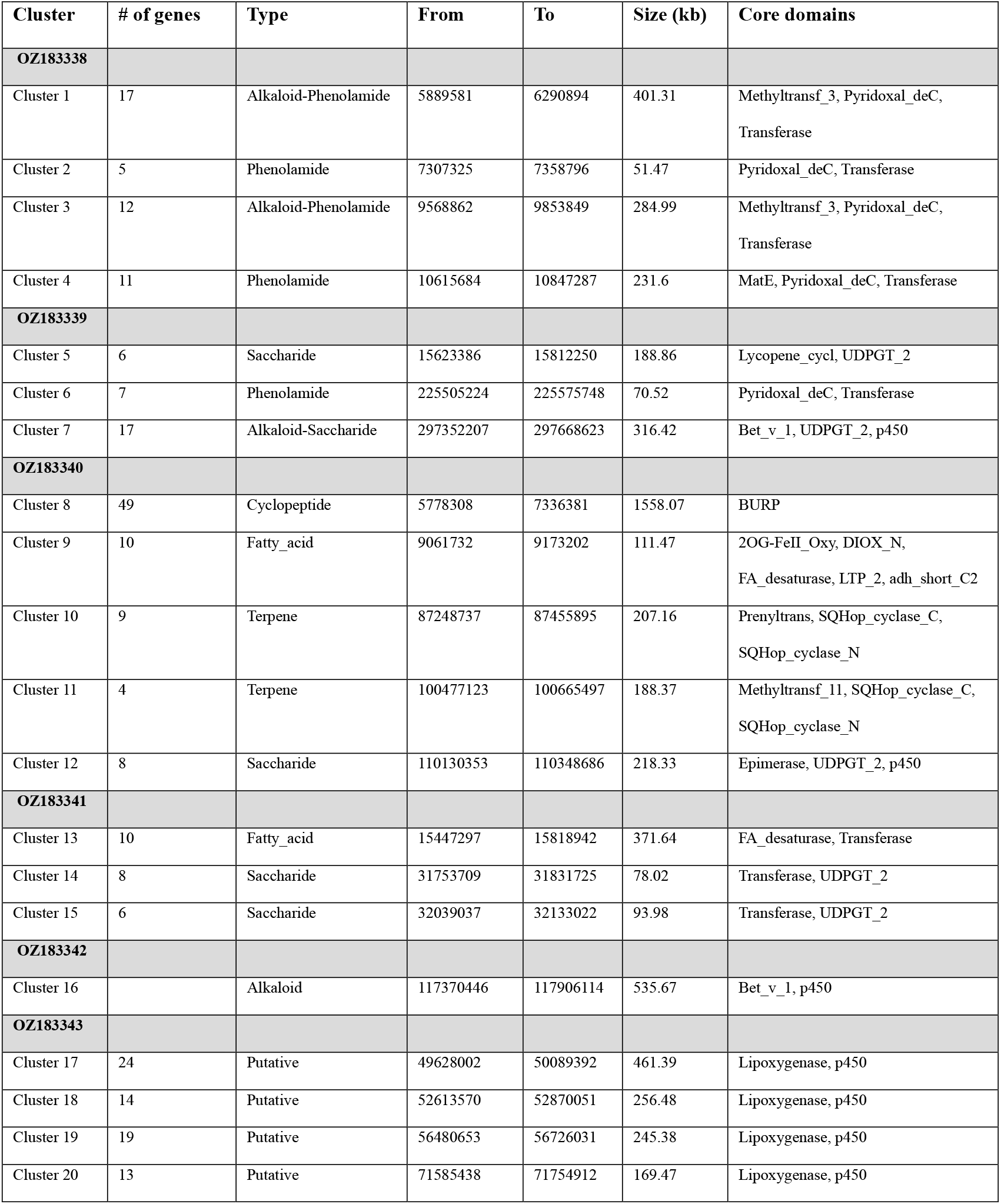
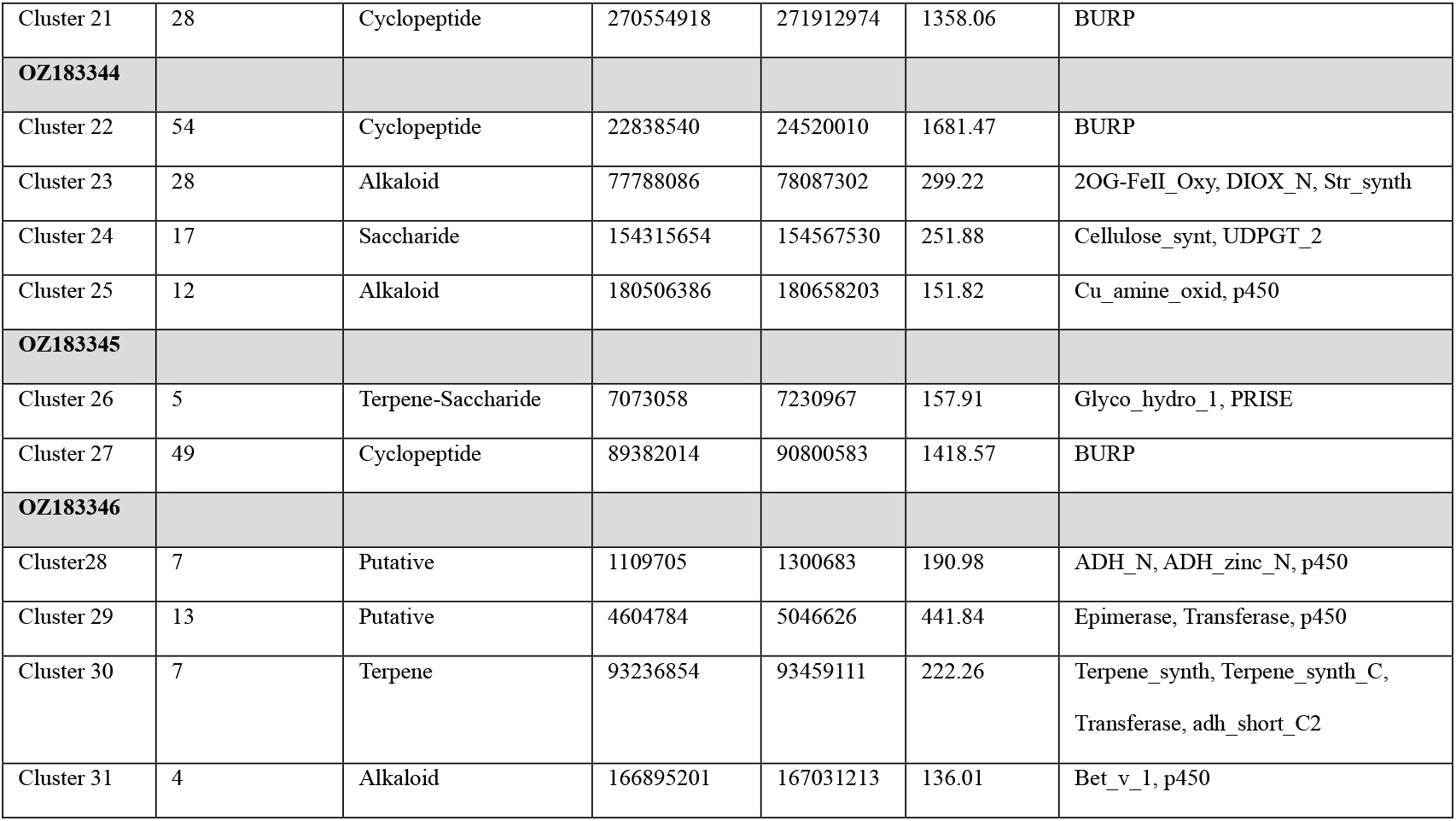
Predicted secondary biosynthetic gene clusters in *Tanacetum vulgare*.

## Data records

The raw sequencing reads generated from all libraries have been deposited in the Ag Data Commons repository. These data are publicly accessible and can be retrieved through the Figshare platform (https://doi.org/10.15482/USDA.ADC/33428869). The archived dataset includes all original FASTQ files and associated metadata required to support reproducibility and downstream analyses.

## Code Availability

No custom scripts were employed in this study. All bioinformatic analyses were conducted in accordance with established protocols and strictly followed the standard command-line instructions provided in the official documentation of each software package.

## Acknowledgments

We thank Jami L. Thomas for optimizing HMW *T. vulgare* gDNA extractions and Brewster F. Kingham and Olga Shevchenko at the University of Delaware DNA Sequencing & Genotyping Center for their valued assistance. Funding for this research was supported by the United States Department of Agriculture, Agricultural Research Service appropriated project 8044-22000-047-00D. AEP and PPE are supported by USDA-HATCH (MICL02742) and MSU AgBioResearch. This research used resources provided by the SCINet project and the AI Center of Excellence of the USDA Agricultural Research Service. The mention of trade names or commercial products in this publication is solely for the purpose of providing specific information and does not imply recommendation or endorsement by the U.S. Department of Agriculture. USDA is an equal opportunity provider and employer.

